# Suppression of miR-155 attenuates lung cytokine storm induced by SARS-CoV-2 infection in human ACE2-transgenic mice

**DOI:** 10.1101/2020.12.17.423130

**Authors:** Dharmendra Kumar Soni, Juan Cabrera-Luque, Swagata Kar, Chaitali Sen, Joseph Devaney, Roopa Biswas

## Abstract

Coronavirus disease 2019 (COVID-19) is a recent global pandemic. It is a deadly human viral disease, caused by the severe acute respiratory syndrome coronavirus 2 (SARS-CoV-2), with a high rate of infection, morbidity and mortality. Therefore, there is a great urgency to develop new therapies to control, treat and prevent this disease. Endogenous microRNAs (miRNAs, miRs) of the viral host are key molecules in preventing viral entry and replication, and building an antiviral cellular defense. Here, we have analyzed the role of miR-155, one of the most powerful drivers of host antiviral responses including immune and inflammatory responses, in the pathogenicity of SARS-CoV-2 infection. Subsequently, we have analyzed the potency of anti-miR-155 therapy in a COVID-19 mouse model (mice transgenic for human angiotensin I-converting enzyme 2 receptor (tg-mice hACE2)). We report for the first time that miR-155 expression is elevated in COVID-19 patients. Further, our data indicate that the viral load as well as miR-155 levels are higher in male relative to female patients. Moreover, we find that the delivery of anti-miR-155 to SARS-CoV-2-infected tg-mice hACE2 effectively suppresses miR-155 expression, and leads to improved survival and clinical scores. Importantly, anti-miR-155-treated tg-mice hACE2 infected with SARS-CoV-2 not only exhibit reduced levels of pro-inflammatory cytokines, but also have increased anti-viral and anti-inflammatory cytokine responses in the lungs. Thus, our study suggests anti-miR-155 as a novel therapy for mitigating the lung cytokine storm induced by SARS-CoV-2 infection.

## INTRODUCTION

Coronavirus disease 2019 (COVID-19) is a life-threatening human viral disease, caused by the severe acute respiratory syndrome coronavirus 2 (SARS-CoV-2) and accountable for the high rate of infection, hospitalization, morbidity and mortality around the world. The outbreak of this deadly human viral pathogen was first reported in December 2019 in Wuhan city of China (1, 2). Subsequently, due to its exceedingly high rate of human-to-human transmission and infection, SARS-CoV-2 rapidly transmitted throughout the world with a very high prevalence, leading to a global pandemic. As of December 8, 2020, almost all (177) countries/regions have reported more than 67,781,393 cases of patients infected with SARS-CoV-2, and 1,548,516 deaths, worldwide, and this continues to rise at an exponential rate (https://coronavirus.jhu.edu/map.html) (3). Therefore, there is great urgency to curb the spread of SARS-CoV-2 and treat infected patients, by developing new therapies to control, treat and prevent this disease. To date, a diverse array of therapeutics COVID-19 is in phase three clinical trial to, such as two mRNA-based vaccine candidate, mRNA-1273 (NIAID and Moderna, Inc.) and BNT162b2 (Pfizer and BioNTech), and a vaccine developed against the SARS-CoV-2 virus spike protein, AZD1222 (University of Oxford and AstraZeneca). Despite the rapid pace with which researchers and manufacturers are moving potential therapeutics into clinical trials, developing effective countermeasures against COVID-19 remains an unmet goal.

The upper respiratory tract is the main site for active replication of SARS-CoV-2, and the severity of the disease depends upon the expression of the viral receptor by lung epithelial cells (4–6). Further, increased inflammatory responses and cytokine storm (such as interleukin (IL)-1β IL-2, IL-6, IL-7, IL-8, tumor necrosis factor-α (TNF-α), granulocyte-colony stimulating factor (G-CSF) and granulocyte-macrophage colony-stimulating factor (GM-CSF)) have been reported in the lungs of severe COVID-19 patients (7–9). In addition, elevated expression of numerous chemokines (monocyte chemoattractant protein 1 (MCP1), interferon-γ inducible protein 10 (IP10) and macrophage inflammatory protein 1-α (MIP1-α)) have also been reported in COVID-19 patients (7, 9, 10). These investigations clearly indicate that the expression of viral specific-receptor by lung epithelial cells have a key role in the entry of SARS-CoV-2 and during infection, immune responses have a critical function in the regulation of inflammation. Therefore, preventing virus entry or replication and/or reinstating or modulating host immunity responses can open a new avenue to control, treat and prevent COVID-19.

Recently, microRNAs (miRNAs, miRs), short (~22 nucleotides) endogenous non-coding RNAs, have emerged as prominent regulators of human defense mechanisms in airway inflammation, lung diseases as well as in various acute respiratory distress syndromes (ARDS). Importantly, in viral infection and disease, the host’s endogenous miRNAs have a decisive function in preventing viral entry and replication either by direct degradation of viral RNA or suppression of translation of proteins involved in replication (11, 12). Furthermore, miRNAs play a predominant role in building antiviral cellular defense and are critical regulators of host immunity and inflammatory responses by inducing mRNA degradation and/or repressing mRNA translation of target genes (13, 14). Over 60% of human genes are targets of miRNAs and are regulated by miRNAs (15, 16). Altered expression of specific miRNAs have been demonstrated in a variety of human diseases including cancer, heart disease, immunological disorders, and diabetes (17–22). Notably, studies have illustrated the association of several miRNAs with immune response against various respiratory viruses such as influenza virus (IV), respiratory syncytial virus (RSV), human rhinovirus (hRV), human metapneumovirus (hMPV), and human coronavirus (HcoV) (12, 23, 24). Some miRNAs regulate levels of immunomodulating factors and suppress or promote inflammatory responses (25). Moreover, since a single miRNA targets multiple genes and modulates the expression of these genes, the best way to nullify the consequence of a dysregulated miRNA in diseased cells, is to restore or inhibit expression (26). Single miRNAs have been shown to stimulate therapeutic responses in animal disease models as well as in human clinical trials (26–29). Therefore, miRNA replacement or inhibitor-based therapeutic mechanisms can be exploited to prevent viral entry or replication and/or restore or modulate host immune responses against COVID-19.

MiR-155, a well-studied miRNA, is one of the most powerful regulators of host antiviral responses including immune and inflammatory responses (30–33). In recent years, numerous studies have clearly established that miR-155 is evolutionarily conserved and during viral infections, its expression is consistently elevated in different cell systems in both human and animal models (34–39). Furthermore, the role of elevated miR-155 is also correlated with lung inflammation, severity of disease and higher mortality in animal models of ARDS (40–43). Interestingly, suppression of miR-155 reduces lung inflammation and induce more rapid recovery of mice infected with respiratory viral pathogens (42, 43). Likewise, knock out (KO) of miR-155 or treatment with anti-miR-155 suppresses cigarette smoke-induced lung inflammation in mice models (44). These studies emphasize the role of miR-155 in the modulation of host defense responses during respiratory virus infection.

Here, we have analyzed the viral load of SARS-CoV-2 and expression level of miR-155 in nasopharyngeal samples of COVID-19 patients. Further, we have evaluated the effect of anti-miR-155 treatment on SARS-CoV-2-infected COVID-19 mouse model, mice transgenic for human angiotensin I-converting enzyme 2 receptor (tg-mice hACE2, tg-mice). Our data indicate that miR-155 expression is significantly elevated in COVID-19 patients. Interestingly, a relatively higher SARS-CoV-2 viral load and miR-155 expression are observed in male COVID-19 patients compared to female patients. Further, our data demonstrate that treatment of SARS-CoV-2-infected tg-mice with anti-miR-155 is effective in suppression of miR-155 expression in the lung tissues, and also promotes improved survival as well as a slight increase in body weight. Moreover, anti-miR-155-treated tg-mice have reduced expression of pro-inflammatory cytokines as well as increased anti-viral and anti-inflammatory cytokines responses in the lungs. These data collectively suggest anti-miR-155 as a novel therapy for mitigating the lung cytokine storm induced by SARS-CoV-2 infection.

## RESULTS

### Higher viral load in SARS-CoV-2 positive male patients

In order to determine the SARS-CoV-2 viral load in COVID-19 patients, a total of 99 human subjects were examined using nasopharyngeal swabs. The isolated RNA was analyzed using COVID-19 RT-PCR diagnostic kit with probes specific for SARS-CoV-2 genes encoded by the RNA-dependent RNA polymerase (RdRp/ORF1ab), Nucleocapsid (N) and Spike (S) protein. Expression of MS2 phage was used as an internal control. From a total of 99 human subjects, 59 (21 male and 38 female; median age, 33 years; range, 12 to 70) tested positive for SARS-CoV-2 infection and 40 (18 male and 22 female; median age, 30 years; range, 13 to 84) tested negative for SARS-CoV-2 infection. Our data indicate that the mean Ct±SEM value for ORF1ab, N and S gene were 21.83±0.84, 23.84±0.84 and 21.36±0.90, respectively, in COVID-19 positive patients (**Fig. 1A**). Further, in the gender-wise grouping analyses, lower Ct values of SARS-CoV-2 genes was observed in male (n=21) relative to female (n=38) patients (**Fig. 1B**), indicating that SARS-CoV-2 viral load is higher in male compared to female COVID-19 patients. However, we acknowledge that larger sample size is required to establish these observations.

**Fig. 1.**
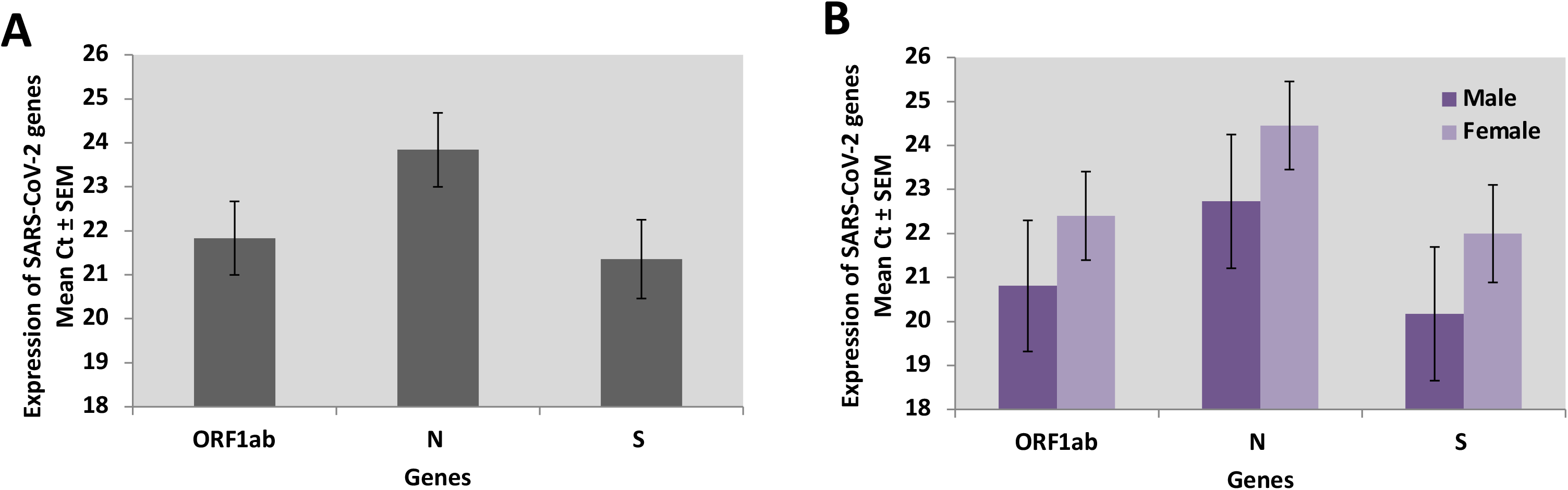
SARS-CoV-2 viral load in COVID-19 patients. The RNA from nasopharyngeal swabs of human subjects (n=99) tested for SARS-CoV-2 infection were analyzed for the expression of SARS-CoV-2 genes, ORF1ab, N and S, by qPCR. The plots depict mean Ct values of the 3 indicated genes analyzed in SARS-CoV-2 positive patients: **(A)** all patients (n=59) and **(B)** male (n=21) *versus* female patients (n=38). The error bars represent ±SEM.

### Elevated miR-155 expression in SARS-CoV-2 positive patients

We analyzed the expression level of miR-155 in the RNA isolated from nasopharyngeal swabs of 99 human subjects (59 positive and 40 negative for SARS-CoV-2 infection) examined for SARS-CoV-2. We find that miR-155 is significantly elevated in patients who tested positive relative to those who tested negative for SARS-CoV-2 infection. Additionally, 34% (n=20) of the patients positive for SARS-CoV-2 infection showed ≥ 2-fold upregulation of miR-155 relative to those who tested negative for SARS-CoV-2 infection (**Figs. 2A and 2B**). Further, in the gender-wise grouping analyses, the level of miR-155 expression is higher in male (n=21, positive *versus* n=18 negative for SARS-CoV-2 infection) relative to female (n=38, positive *versus* n=22, negative for SARS-CoV-2 infection) patients (**Fig. 2C**). These results indicate that miR-155 is a potential target for COVID-19 therapy.

**Fig. 2.**
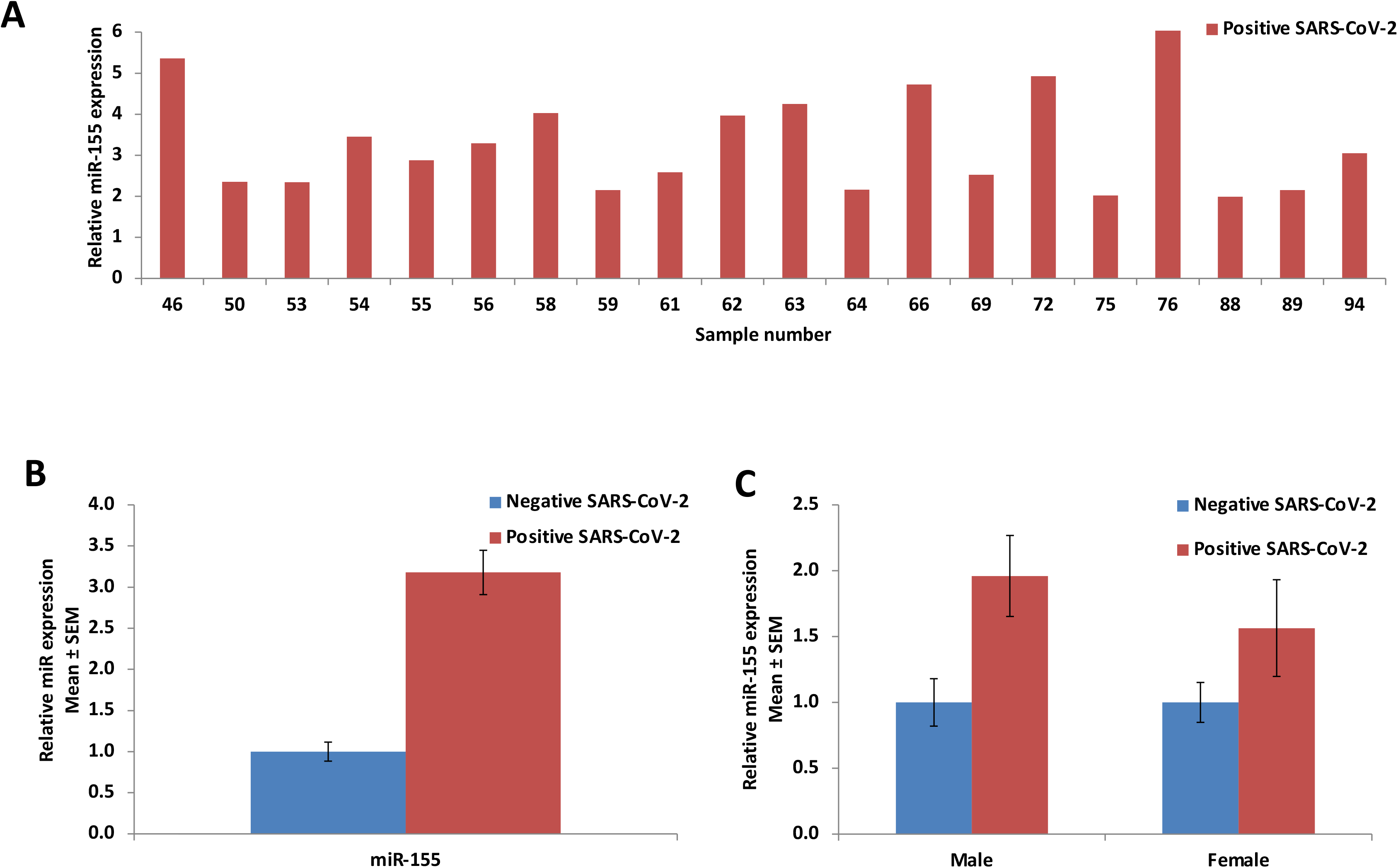
Expression of miR-155 in SARS-CoV-2 positive patients. The expression of miR-155 in **(A)** individual SARS-CoV-2 positive patients and **(B)** average levels in SARS-CoV-2 positive patients with ≥ 2 fold upregulation (n=20) compared to SARS-CoV-2 negative (n=40) was analyzed by Taqman assay. **(C)** The relative miR-155 levels in male patients (positive n=21 *versus* negative n=18 for SARS=CoV-2) and female patients (positive n=38 *versus* negative n=22 for SARS=CoV-2) are depicted.

### Delivery of anti-miR-155 suppressed miR-155 expression in lung tissues of SARS-CoV-2-infected tg-mice hACE2

We evaluated the efficacy of *in vivo* delivery of anti-miR-155 to the lungs of SARS-CoV-2-infected tg-mice hACE2 (strain 034860 B6.Cg-TG (K18-ACE2)2Prlmn/J), obtained from Jackson Laboratory (Bar Harbor, ME, USA). Tg-mice hACE2 (8-10-week-old) were infected via intranasal injection with 2.8 × 10^3^ plaque-forming units (pfu) of SARS-CoV-2 strain USA-WA1/2020 (BEI Resources NR-52281, batch #70033175). Subsequently, the tg-mice were treated with anti-miR-155 or anti-miR-control (scrambled control) via retro-orbital injection at a dose of 25 mg/kg body weight (BW) in 50 μL PBS on 0-and 4-days post-infection (dpi). Total RNA was isolated from the lungs of tg-mice and the expression of miR-155 was measured using miR-155-specifc Taqman assay. As depicted in **Fig. 3A,** a significant suppression of miR-155 (~30%) is observed in the lung tissues of SARS-CoV-2-infected anti-miR-155-treated tg-mice (n=6) compared to anti-miR-control tg-mice (n=3). These results demonstrate the effectiveness of anti-miR-155, administered via retro-orbital injection, in suppressing miR-155 in the lung tissues of SARS-CoV-2-infected tg-mice. The dose and administration regime will have to be further optimized.

**Fig. 3.**
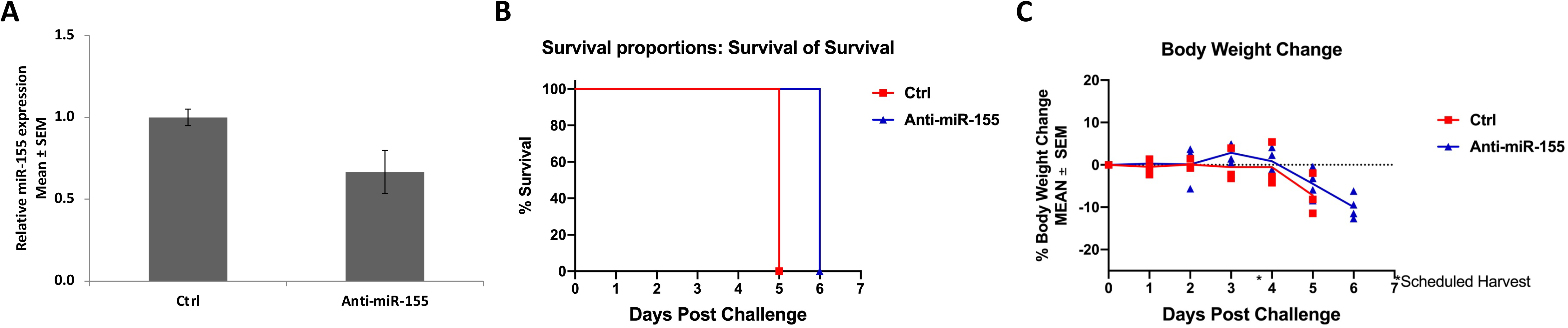
Impact of anti-miR-155 treatment on tg-mice hACE2 infected with SARS-CoV-2. Tg-mice hACE2 was infected by intranasal injection with SARS-CoV-2 at 2.8 × 10^3^ pfu, were treated with anti-miR-155 (n=6) or anti-miR-control (Ctrl, scrambled, n=3) at a dose of 25 mg/kg mice body weight administered on 0- and 4-day post-infection via retro-orbital injection. **(A)** Expression level of miR-155 in lung tissues of anti-miR-155-treated tg-mice (n=6) compared to control tg-mice (n=3) was measured. **(B)** Kaplan-Meier survival curve depicts the differential survival of anti-miR-155-treated compared to control tg-mice. **(C)** Percent body weight change in the two groups of animals is shown.

### Improved survival and body weight of SARS-CoV-2-infected tg-mice after treatment with anti-miR-155

We examined the effect of anti-miR-155 treatment on BW and survival of tg-mice hACE2 infected with SARS-CoV-2. The anti-miR-155-treated infected tg-mice (n=6) had relatively higher survival compared to the anti-miR-control-treated mice (n=3) (**Fig. 3B**). All control mice succumbed to infection and were euthanized on 5 dpi, while the anti-miR-treated tg-mice survived a day longer and succumbed to infection and were euthanized on 6 dpi. Further, we observed that the anti-miR-155-treated tg-mice showed a slight increase in or stable BW (with a peak during days 2-4), compared to control tg-mice (**Fig. 3C**). However, subsequently the animals from both anti-miR-155-treated and control group developed clinical signs of disease (weight loss, mild ruffled and hunched back, lethargic, not alert and difficulty breathing). From 4 dpi onwards, SARS-CoV-2-infected anti-miR-155-treated tg-mice as well as control mice displayed a rapid reduction in BW (**Fig. 3C**). These results emphasize the role of anti-miR-155-based therapeutics for COVID-19 and suggests that perhaps a more frequent administration regime, i.e., treatment on alternate days post-infection, could improve the therapeutic efficacy.

### Altered levels of inflammatory mediators in the lungs of SARS-CoV-2-infected tg-mice treated with anti-miR-155

We evaluated the responses of anti-miR-155 treatment on cytokine and chemokine profile in the lungs of SARS-CoV-2-infected tg-mice. Comprehensive analyses of cytokine profile (Luminex, 36-plex MILLIPLEX® MAP Mouse Cytokine/ Chemokine Magnetic Bead kit, EMD Millipore) in the bronchoalveolar lavage (BAL) fluid indicate attenuation of the “cytokine storm” in anti-miR-155-treated tg-mice (n=5) compared to control mice (n=1). We were unable to obtain BAL fluid from one of the treated tg-mice and 2 control tg-mice. The data indicate significant downregulation of cytokines and chemokines including G-CSF, IL-1α, IL-9, IL-12 (p70), KC, LIX, MIP-2, and VEGF (**Fig. 4**). Additionally, other cytokines, which are relevant to SARS-CoV-2 infection, such as eotaxin, IFN-γ, IP-10, MCP-1, M-CSF, MIG, MIP-1α, MIP-1β, and TNF-α are induced in anti-miR-155-treated tg-mice compared to control tg-mice (**Fig. 5**). Importantly, a significant elevation of IFN-γ, IP-10, and TNF-α suggests the antiviral potency of anti-miR-155 in addition to its anti-inflammatory efficacy in the tg-mice model of COVID-19. These data collectively suggest the anti-inflammatory as well as the anti-viral potency of anti-miR-155 therapy for COVID-19.

**Fig. 4.**
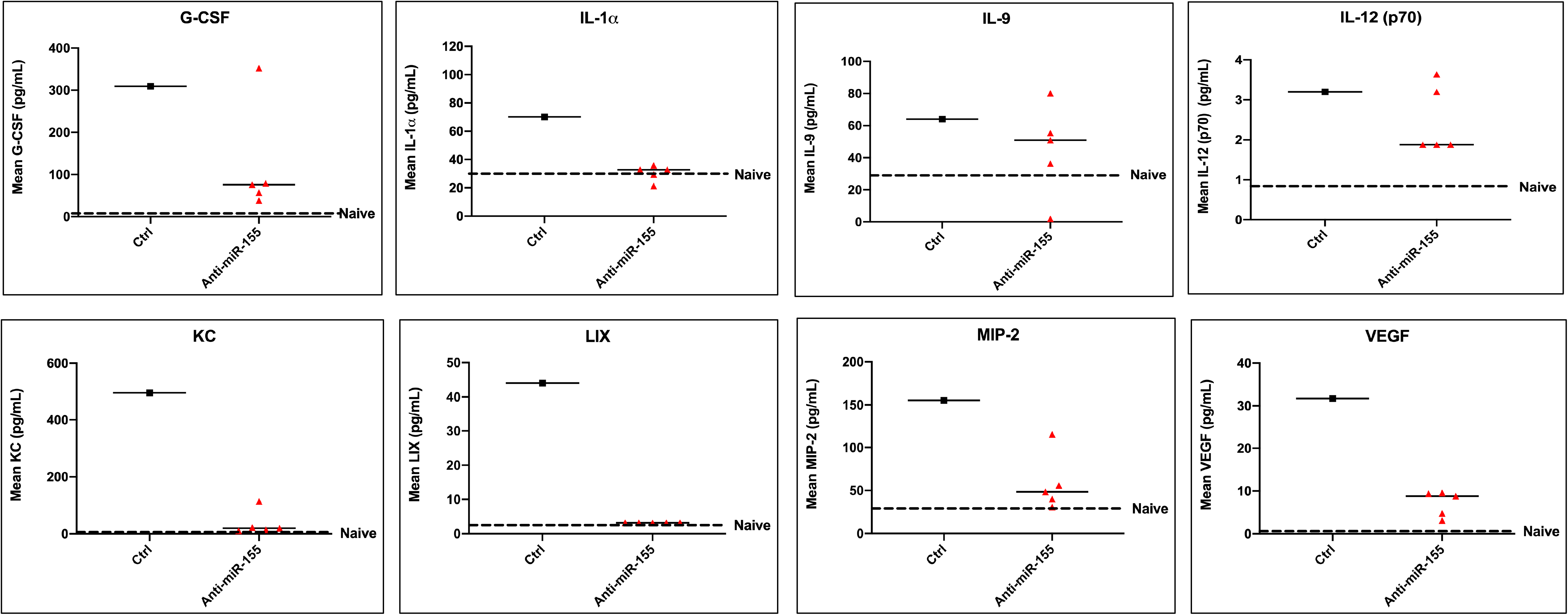
Impact of anti-miR-155 treatment on pro-inflammatory cytokines. The BAL fluid obtained from SARS-CoV-2-infected tg-mice hACE2 treated with anti-miR-155 (n=5), and tg-mice treated similarly with anti-miR-control (Ctrl, scrambled, n=1) was analyzed for cytokine profile by Luminex. Relative levels of the cytokines, G-CSF, IL-1α, IL-9, IL-12 (p70), KC, LIX, MIP-2, and VEGF, significantly altered are shown.

**Fig. 5.**
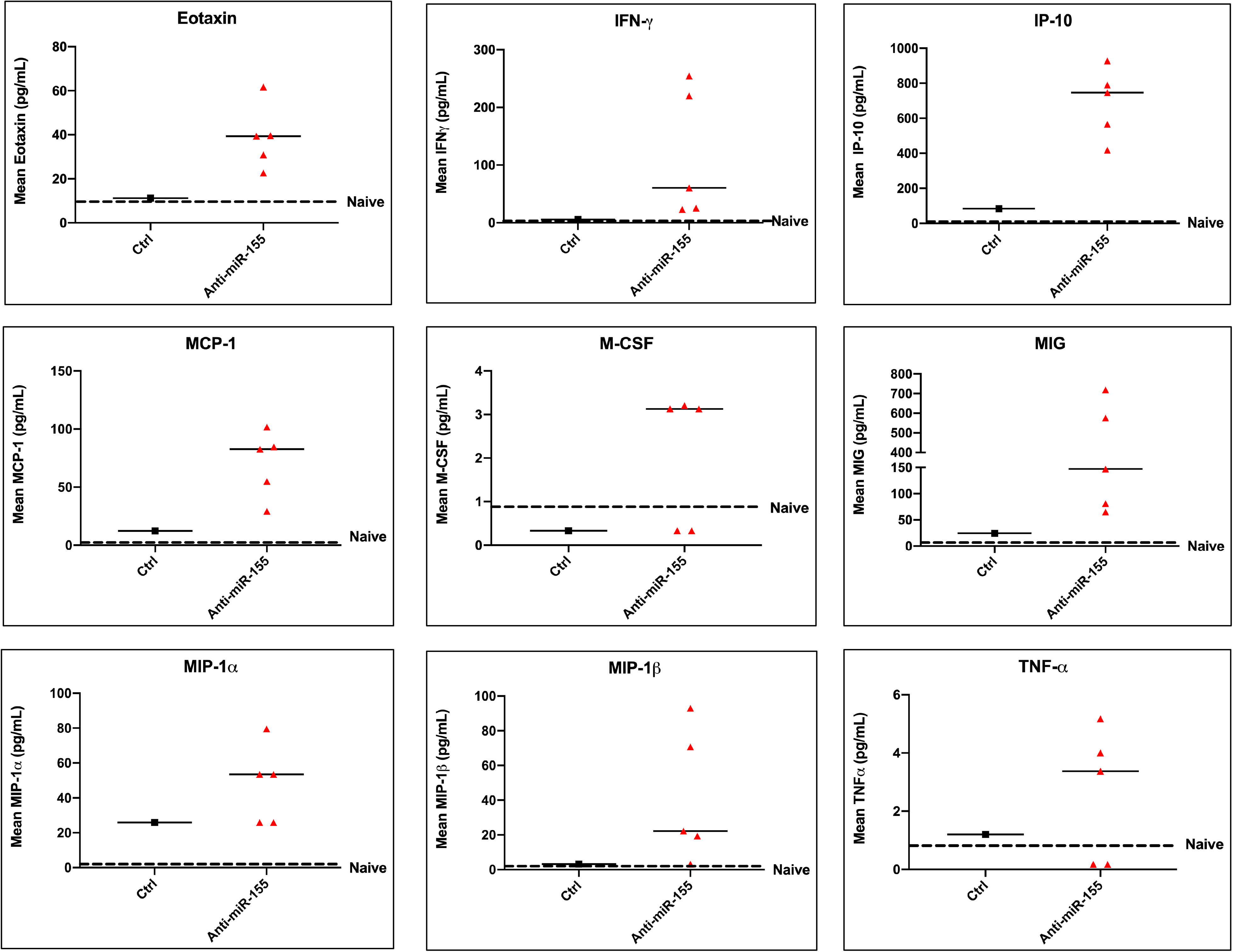
Impact of anti-miR-155 treatment on anti-inflammatory and anti-viral cytokines. The cytokine profile of the BAL fluid obtained from SARS-CoV-2-infected tg-mice hACE2 treated with anti-miR-155 (n=5) or with anti-miR-control (Ctrl, scrambled, n=1) was analyzed by Luminex. Relative levels of the cytokines, eotaxin, IFN-γ, IP-10, MCP-1, M-CSF, MIG, MIP-1α, MIP-1β, and TNF-α, that are significantly altered are shown.

## DISCUSSION

From the beginning of this year 2020, the entire world is facing challenges to human health as well as an economic threat because of COVID-19, a deadly human disease caused by a viral pathogen SARS-CoV-2. This had created a global urgency for developing therapeutics against this life-threatening pathogen to control, treat and prevent COVID-19. Currently, many therapeutics strategies are in phase three clinical trials, such as mRNA-1273, BNT162b2, and AZD1222. Despite these recent advances, developing effective countermeasures against this lethal pathogen remains an unmet goal.

SARS-CoV-2 infection primarily affects the lungs and exacerbates host immune and inflammatory responses, which is one of predominant cause for increased cellular damage, severity of disease and death (45, 46). Several studies have illustrated the potential of miRNAs, including miR-155 in the regulation of antiviral cellular defense mechanisms and mitigation of infections caused by viral pathogens (30–33, 43, 47–53). However, to date there are no reports of the impact of miR-155 on SARS-CoV-2 infections. Here, we investigated the expression of miR-155 in the nasopharyngeal samples of COVID-19 patients. Our data indicate significant upregulation of miR-155 expression in SARS-CoV-2 positive patients compared to those who tested negative. Interestingly, we observed higher SARS-CoV-2 viral load and higher upregulation of miR-155 in SARS-CoV-2 positive male relative to the female patients. However, to establish the gender effect observed analyses of a larger number of patients is required. Consistent with our data, miR-155 has been shown to be induced in the SARS-CoV-2-infected Calu-3 lung cell line (54). Another report indicates that only two miRNAs, including miR-155-3p and miR-139-5p, are upregulated in lung epithelial cells upon SARS-CoV-2-infection with a significant prediction score (55). Furthermore, recent studies have clearly established that miR-155 is evolutionarily-conserved and during viral infections, its expression is consistently elevated in animal and human models (34–39). Our findings, significantly elevated expression of miR-155 in COVID-19 patients, strongly suggest the regulatory role of miR-155 during SARS-CoV-2 infection.

Interestingly, the elevated expression of miR-155 has been correlated with the severity of disease in ARDS. To define the role of miR-155 during SARS-CoV-2 pathogenesis, we further evaluated the effect of anti-miR-155 treatment on SARS-CoV-2-infected tg-mice hACE2. The wild-type mice are not suitable animal models for studying SARS-CoV-2 infection or the resulting pathologies because the virus does not bind as effectively to the murine ACE2 protein receptor as to ACE2 receptors in humans and several other species. However, tg-mice hACE2 develops severe disease following infection with beta-coronaviruses, including SARS-CoV-2 (56–58). Our data demonstrate significant suppression of miR-155 expression in the lungs of SARS-CoV-2-infected anti-miR-155-treated tg-mice relative to control tg-mice, which establishes the *in vivo* delivery efficacy as well as the effectiveness of anti-miR-155 treatment (administered via retro-orbital injection). Further, the anti-miR-155-treated tg-mice exhibited a slight increase in or stable BW compare to the control tg-mice. However, subsequently both anti-miR-155-treated and control groups of animals developed clinical signs of disease (weight loss, mild ruffled and hunched back, lethargic, not alert and difficulty breathing). From 4 dpi, both SARS-CoV-2-infected anti-miR-155-treated and control tg-mice displayed a rapid reduction in BW. Our findings are consistent with previous studies which have reported low survival and significant weight loss in SARS-CoV-2-infected tg-mice (58, 59). Additionally, our results demonstrated that SARS-CoV-2-infected anti-miR-155-treated tg-mice showed slightly improved survival compared to control tg-mice. Studies on viral infections associated with ARDS have reported that the elevated expression of miR-155 in whole lung tissue is linked to higher mortality and severity of disease in mice (40–43). Furthermore, deletion or suppression of miR-155 has been also correlated with decreased lung inflammation and increased recovery of mice infected with respiratory viral pathogens (42, 43). The impact of anti-miR-155 treatment in promoting increased survival and BW of mice suggest a vital role of miR-155 in the pathogenesis of SARS-CoV-2. Further, our data also indicate that perhaps a more frequent administration regime, i.e., treatment on alternate days post-infection, could improve the therapeutic efficacy of anti-miR-155.

The high mortality rate of SARS-CoV-2-infected patients has been associated with aggressive inflammatory responses and exaggerated immune responses termed as a cytokine storm. The cytokine storm is characterized by excessive production of various pro-inflammatory cytokines and chemokines, such as TNF-α, IL-1β, IL-2, IL-6, IL-7, IL-8, G-CSF, GM-CSF, MCP1, IP10 and MIP1α) (7, 9, 10, 46, 60–63). The role of miR-155 is well-established in the induction of pro-inflammatory cytokines and chemokines during numerous viral diseases including ARDS. Therefore, we hypothesized that suppression of miR-155 expression could attenuate excessive production of pro-inflammatory cytokines and chemokines and thereby, mitigate the lung cytokine storm induced by SARS-CoV-2-infection. With this hypothesis, we further evaluated the responses of anti-miR-155 treatment on cytokine and chemokine expression profile in the lungs of SARS-CoV-2-infected tg-mice. The tg-mice infected with SARS-CoV-2 exhibits increased pro-inflammatory responses as observed in COVID-19 patients (57, 58). The tg-mice infected with SARS-CoV-2 has increased serum protein levels of TNF-α, IL-6, and IL-10 as well as the MCP-1 (CCL2) and MCP-3 (CCL7) in the lung homogenates (57). Additionally, the SARS-CoV-2-infected tg-mice also have increased protein levels of interferon-β (IFN-β), IFN-γ, IL-2, IL-6, IL-10, C–X–C motif chemokine ligand 1 (CXCL1), CXCL9, CXCL10, chemokine ligand 2 (CCL2), CCL3, CCL4, CCL5, CCL12, tissue inhibitor of metalloproteinase 1, TNF-α and G-CSF in the lungs (58). Our data indicate that in SARS-CoV-2-infected anti-miR-155-treated tg-mice, BALF inflammatory mediator (cytokines and chemokines) levels are altered with the decrease in G-CSF, IL-1α, IL-9, IL-12 (p70), KC, LIX, MIP-2, and VEGF and increase in eotaxin, IFN-γ, IP-10, MCP-1, M-CSF, MIG, MIP-1α, MIP-1β, and TNF-α. Importantly, we observed a significant elevation of IFN-γ, IP-10, and TNF-α, suggesting the antiviral potency of anti-miR-155 in addition to its anti-inflammatory efficacy in the tg-mice model of COVID-19. Previously, Golden et al. also reported the decreased expressions of Type I and type II IFN transcripts IFNA1, IFNB1, and IFNG, along with the cytokines IL-9 and IL-2 in the lung homogenates of tg-mice upon SARS-CoV-2 infection (57). Similarly, Winkler et al. also reported the variation in the expressions of various cytokines and chemokines in the lungs of SARS-CoV-2-infected tg-mice, while some are increased, some showed no change, and others rapidly increased or decreased following infection (58).

## CONCLUSION

MiR-155 is elevated in the nasopharyngeal samples of COVID-19 patients. The *in vivo* suppression of miR-155 in SARS-CoV-2-infected tg-mice hACE2 promotes improved survival and can mitigate the cytokine storm and lung inflammation. Thus, our study suggests anti-miR-155 as a novel therapeutic candidate for COVID-19.

## METHODS

### Detection of SARS-CoV-2 and measurement of viral load in human patients

Ninety-nine human subjects (39 male and 60 female; median age, 32 years; range, 12 to 84), who were examined for SARS-CoV-2 infection at OPKO Health Company (GeneDx, Inc., MD, USA) were included in this study. Nasopharyngeal swab samples were collected in 3mL of standard viral transportation media (VTM). Tubes containing the swab and VTM were pulse-vortexed three times and 1.5mL aliquots were collected onto low protein binding tubes (ThermoFisher cat# 88379). Tubes were then centrifuged at 4⁰C for 10 minutes at 1500g. VTM was carefully removed from the tubes and RNA was extracted from pelleted cells. The detection and measurement of SARS-CoV-2 viral load were performed using COVID-19 RT-PCR diagnostic kit with specific Taqman probes (ThermoFisher) of SARS-CoV-2-specific genes encoded by the RNA-dependent RNA polymerase (RdRp/ORF1ab), Nucleocapsid (N) and spike (S) protein. MS2 phage was used as an internal control. The expression of the SARS-CoV-2 genes was analyzed by the corresponding Ct values, which are inversely proportional to the viral load.

### Measurement of miR-155 expression in human patients

For analyses of miRNA expression, the RNA was extracted from the VTM using the RNAqueous Micro Kit (ThermoFisher), following manufacturer specifications. During RNA isolation, 100fmol of cel-miR-39 RNA (Ambion) was added as spike-in-control. The expression of miR-155 and cel-miR-39 were analyzed by specific Taqman assays (ThermoFisher). Cel-miR-39 was used as an endogenous control to normalize expression. The relative fold changes of miR-155 were analyzed using the 2^−ΔΔCT^ method.

### Mouse model

Eight to ten weeks old mice transgenic for human angiotensin I- converting enzyme 2 receptor (tg-mice hACE2, tg-mice; average weight =22 gm) strain 034860 B6.Cg-TG(K18-ACE2)2Prlmn/J were obtained from Jackson Laboratory (Bar Harbor, ME, USA). Animals were housed in groups of five animals per cage and fed standard chow diets. All the experiments with live virus challenge were carried out at BIOQUAL Inc. (Rockville, MD, USA) bio-safety level 3 (BSL-3) facilities in compliance with local, state, and federal regulations under IACUC protocol #20-083.

### Virus challenge

SARS-CoV-2 strain USA-WA1/2020 (BEI Resources NR-52281, batch #70033175), courtesy of the Centers for Diseases Control and Prevention (CDC), was used for mice infection. The SARS-COV-2 stock was expanded in Vero E6 cells, and the challenging virus was collected at day 5 of culture when the infection reached 90% cytopathic effect. The full viral genome sequence showed 100% identity with the parent virus sequence listed in GenBank (MN985325.1). A plaque-forming assay was carried out in confluent layers of Vero E6 cells and used to determine the concentration of live virions, reported as plaque-forming units (pfu). Virus challenge was performed in tg-mice hACE2 via intranasal injection with 2.8 × 10^3^ pfu of SARS-CoV-2. Intranasal administration was performed by anesthetizing the animal with Ketamine/ Xylazine. Animals are returned to their housing unit and monitored until fully recovered from the procedure. Bodyweight and clinical observations were monitored daily during the experimental period.

### Administration regime for anti-miR-155

SARS-CoV-2-infected tg -mice hACE2 were treated with anti-miR-155 at a dose of 25 mg /kg (for an average weight of 22 g per mouse) in 50 μL PBS at 0-and 4-days post-infection (dpi) via Retro-Orbital (RO) injections. A set of control infected tg-mice (n=3) were treated similarly with anti-miR (scrambled)-control. Following the infection of tg-mice with SARS-CoV-2, the lungs from the set of control tg-mice treated with anti-miR (scrambled)-control were isolated on 5 dpi, and the lungs from the anti-miR-155-treated tg-mice were isolated on 6 dpi. All isolated lungs from control and treated tg-mice were immediately snap-frozen over dry ice and stored at −80°C.

### Measurement of miR-155 expression in animals

The total RNA was isolated from frozen lung tissues using TriZol reagent (Invitrogen Inc., Carlsbad, CA) with a *mir*Vana™ miRNA isolation kit (Ambion, AM1561) and quantitated using Nanodrop (Thermo Scientific). MiRNAs analysis was performed with specific Taqman probes and U6 was used as an endogenous control to normalize expression of miR-155. The relative fold changes of miR-155 were analyzed using the 2^−ΔΔCT^ method.

### Measurement of cytokine and chemokines

The level of cytokine and chemokines protein in BAL samples from anti-miR-155-treated tg-mice hACE2 and anti-miR (scrambled)-control tg-mice hACE2 were analyzed by Luminex multiplex assays using 36-plex MILLIPLEX® MAP Mouse Cytokine / Chemokine Magnetic Bead kit, (EMD Millipore) according to the manufacturer’s protocol.

### Statistical analysis

Statistical analysis for viral load and miR-155 expression was performed using Excel. The data represents average Ct values or fold changes with the standard error of the mean (mean ± SEM). The percent change in body weight, Kaplan-Meier survival analyses and alteration in cytokine expression in the experimental compared to control mice were analyzed with Prism version 8 (GraphPad).

## FOOTNOTES

### Disclaimer

The views expressed are those of the authors and do not reflect the official policy or position of the Uniformed Services University of the Health Sciences, the Department of the Defense, or the United States government.

## Abbreviations used

COVID-19: coronavirus disease-19
SARS-CoV-2: severe acute respiratory syndrome coronavirus 2
hACE2: human angiotensin I-converting enzyme 2 receptor
miRNA: miR, microRNA
BAL: bronchoalveolar lavage.

## Acknowledgements

The authors thank Samarjit Das (Johns Hopkins University) for helpful discussions, Maciel Porto (Bioqual Inc.) for monitoring the animal study and performing Luminex assays, and Melissa Hamilton (Bioqual Inc.) for processing and analyses of data.

## Funds

This study was supported by NHLBI/USUHS-CHIRP funds 1I80VP000012 and USUHS Intramural Grant [RB].

